# Perception of soft materials relies on physics-based object representations: Behavioral and computational evidence

**DOI:** 10.1101/2021.05.12.443806

**Authors:** Wenyan Bi, Aalap D. Shah, Kimberly W. Wong, Brian Scholl, Ilker Yildirim

## Abstract

When encountering objects, we readily perceive not only low-level properties (e.g., color and orientation), but also seemingly higher-level ones – including aspects of physics (e.g., mass). Perhaps nowhere is this contrast more salient than in the perception of soft materials such as cloths: the dynamics of these objects (including how their three-dimensional forms vary) are determined by their physical properties such as stiffness, elasticity, and mass. Here we hypothesize that the perception of cloths and their physical properties must involve not only image statistics, but also abstract object representations that incorporate ”intuitive physics”. We provide behavioral and computational evidence for this hypothesis. We find that humans can visually match the stiffness of cloths with unfamiliar textures from the way they undergo natural transformations (e.g. flapping in the wind) across different scenarios. A computational model that casts cloth perception as mental physics simulation explains important aspects of this behavior.

## Introduction

When encountering objects, we readily perceive not only low-level properties (e.g., color and orientation), but also seemingly higher-level ones – some of which seem to involve aspects of physics (e.g., mass and stiffness). Perhaps nowhere is this contrast more salient than in the perception of soft materials such as cloths: the dynamics of these objects (including how their three-dimensional forms vary) are determined by their physical properties such as stiffness, elasticity, and mass.

Existing work emphasize the role of the statistical regularities in the low- or mid-level image features as clues to the estimation of mechanical material properties (Fleming, 2017; Nishida, Kawabe, Sawayama, & Fukiage, 2018; Nishida, 2019; Van Assen, Barla, & Fleming, 2018). This hypothesis is supported by studies that have found that a variety of image features, such as two-frame motion cues (Kawabe, Maruya, Fleming, & Nishida, 2015; Kawabe & Nishida, 2016), shape deformation (Paulun, Kawabe, Nishida, & Fleming, 2015; Paulun, Schmidt, van Assen, & Fleming, 2017; Schmidt, Paulun, van Assen, & Fleming, 2017; Schmidt, Fleming, & Valsecchi, 2020), and multi-frame motion information (Bi, Jin, Nienborg, & Xiao, 2018), can affect the visual perception of mechanical properties. However, we do not perceive individual material properties just as patterns or statistical regularities, but in the mind they are cast as rich, structured physical representations of entities and soft objects that we can think about and predict future motions of. Consider the veiled sculpture in Figure 1: despite knowing its true material (a marble sculpture), we see a soft object, a veil, with a certain mass and softness that appears to be interacting with the blowing wind. Similarly, in the hanging canvas, despite its unfamiliar texture (from its intricate strokes of paint), we can tell the stiffness of the canvas and estimate its weight from the folds it makes under the influence of gravity. Such perception is more than just patterns or statistical regularities. How are soft objects represented in the mind so as to support such rich inferences that often go far beyond what is in the sense inputs? And how are these representations inferred and updated from sequential, dynamic inputs?

**Figure 1:**
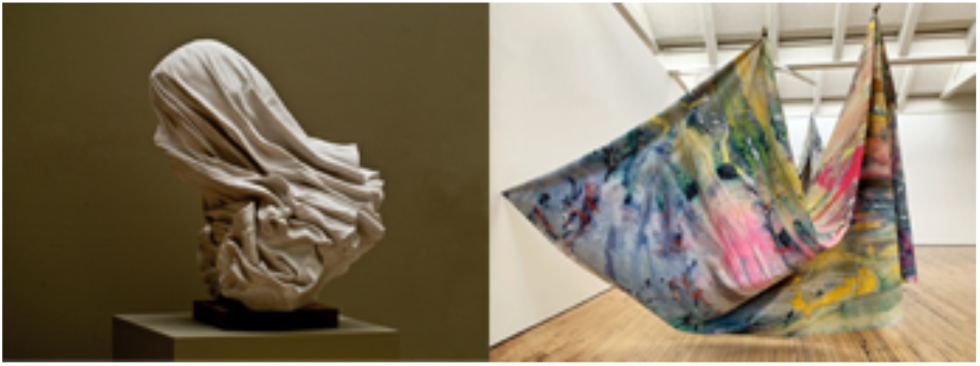
Soft material, e.g. cloth, often induce rich object representations. Left: The Veiled Woman by Rubincam (2013); Right: the canvas of a hanging painting by Davis (2019).

Inspired by the recent work in several domains of physical scene understanding (Battaglia, Hamrick, & Tenenbaum, 2013; C. J. Bates, Yildirim, Tenenbaum, & Battaglia, 2019; Wu, Yildirim, Lim, Freeman, & Tenenbaum, 2015; Yildirim, Siegel, & Tenenbaum, 2016, 2020), here we argue that visually estimating physical properties of cloth must involve not only image statistics, but also abstract object representations that incorporate ”intuitive physics” – an abstract, physics-based representation of approximate cloth mechanics that explains observed shape variations in terms of how unobserved object properties determine cloth reaction to external forces. We realize this hypothesis in a computational model of cloth perception. In this model, perception of soft object properties (e.g., mass, stiffness) is cast as approximate probabilistic inference in a simulation-based generative model. This model incorporates physical principles of how cloth mechanics work and how soft objects react to external forces in real-time, and the efficient simulations that take shortcuts to approximate the otherwise highly complex cloth dynamics. We show that a Sequential Monte Carlo (SMC) algorithm with limited computational resources (very few particles) can make accurate inferences of physical cloth properties.

Additionally, we provide behavioral and computational evidence for our hypothesis by studying the ability to “generalize” across different scenarios and external force types. We do so by evaluating the human behavior and the computational model’s performance on a 2AFC visual matching task. In this task, we asked observers to visually match the stiffness of animated cloths reacting to external forces and undergoing natural transformations (e.g., flapping in the wind, falling onto an uneven surface). In our analysis, we find that the human matching performance is robust despite the massive variability in the lower-level image statistics and the higher-level variability in both extrinsic scene forces (e.g., wind vs. rigid-body collision) and intrinsic cloth properties (e.g., mass). Via computational modeling, we find that our model makes intuitive trade-offs between its inferences about the stiffness and mass of observed cloths, and when these inferences are used to perform the aforementioned visual matching task, it captures important variations in the behavioral error patterns. These results open the domain of the mental representations of soft objects into computational inquiry, and suggest an account based on internal models of their mechanics and how they react to forces.

## Task: Perceptual generalization of cloth physical properties across different scenarios

Most previous studies on the perception of non-rigid objects require participants to compare physical properties (e.g., liquid viscosity, elasticity of jellylike cubes) within the same scenario (Bi & Xiao, 2016; Kawabe & Nishida, 2016; van Assen & Fleming, 2016). Because force type is constant, the brain doesn’t need to decode the physical principles that determine how the object deforms in reaction to the forces—estimating physics is relatively straightforward such that the visual system can detect a fixed set of image features to enable robust estimation of the physical properties.

Comparing physical properties across scenarios is much more challenging—image features vary greatly across scenes, such that the robust image features for estimation of physical properties in one scene might not even exist in another scene. To disentangle the external forces from intrinsic physical properties and to achieve their invariant estimation, the brain needs to decode certain physical principles and perform causal inference based on those principles (e.g., through mental simulation). Thus, being able to generalize across scenarios suggests the involvement of physics-based representations of soft objects when estimating their physical properties.

Here we designed four scenarios containing a cloth reacting to different types of external forces and undergoing natural transformations (Figure 2A). We adopted a 2AFC procedure that requires participants to compare cloth stiffness across scenarios (Figure 2B).

**Figure 2:**
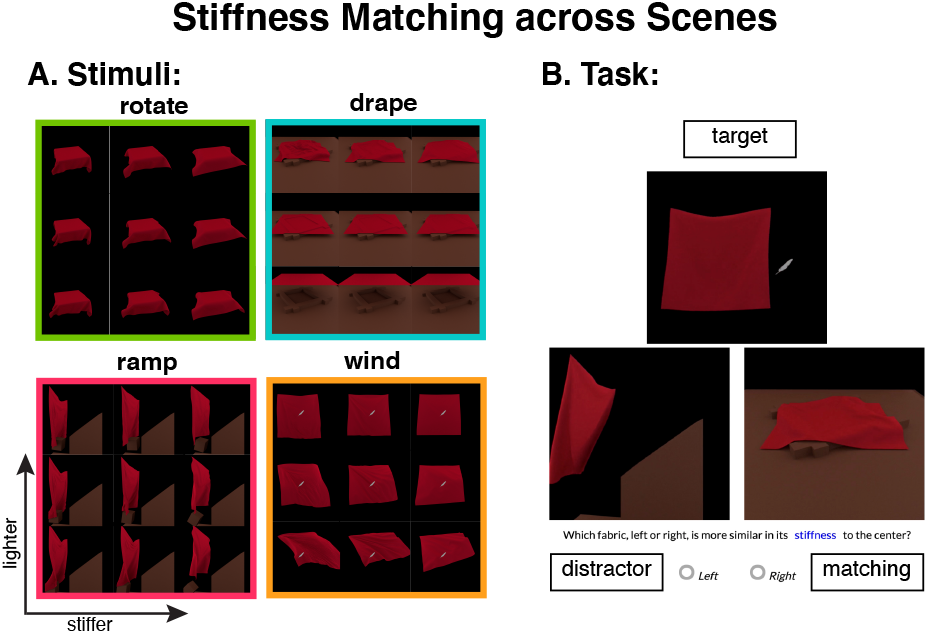
A. Example stimuli used in the experiment. We created 4 different scenarios containing a cloth reacting to different types of external forces and undergoing natural transformations—image statistics are very different across different scenarios. For each of the four scenarios, the nine images were taken from the same time step: the upper three images illustrate the lightest cloth, and the lower three illustrate the heaviest. Similarly, the left three images show the softest cloth, and the right three show the most stiff. B. 2AFC stiffness matching task. On each trial, participants were presented with videos triads from three different scenarios, and were asked to decide which of the two cloths on the bottom had the same stiffness value as that of the one on the top (i.e., target).

Figure 2A shows example frames of the video stimuli that we used. Stimuli consisted of computer-rendered animations of cloth reacting to external forces in four different scenarios: (1) A rotate scenario (upper left panel) consisted of a cloth laid on a rotating table. The table started rotating with a fixed angular velocity after the cloth was fully draped on it. (2) A drape scenario (upper right panel) consisted of a cloth falling under gravity, from a fixed height, onto a wood frame placed on the floor. (3) A ramp scenario (lower left panel) contained a box sliding down a ramp and colliding with a hanging cloth. The initial height, velocity (i.e., 0), and weight of the box was fixed. (4) A wind scenario consisted of a hanging cloth being blown by unknown oscillating winds. The wind profile was fixed and could be appreciated by the movements of a light feather in the scene. The animations were simulated in Nvidia FleX (Macklin, Müller, Chentanez, & Kim, 2014) with the same set of parameters, including the cloth size (105 × 105), friction coefficients, damping coefficients, iterations, time steps, etc. The cloth in each scenario was simulated with a wide range of stiffness (2^−7^,2^−5^,2^−3^,2^−1^,2^1^) and mass (2^−2^,2^−1^,2^0^,2^1^,2^2^) values with a total of 100 unique cloth animations, each having 200 frames. The animations were rendered using Blender (v. 2.7.9.) Cycles Render Engine with the same rendering parameters for the four scenarios except for a slight difference in the camera angle.

Figure 2B shows the 2AFC stiffness matching procedure used in this experiment. On each trial, observers were presented with video triads – one “target” video on the top and two ‘alternate” videos on the bottom – and they were asked to decide which of the two alternative cloths had the same stiffness value compared to the target. Participants indicated their choice by clicking the “left”/”right” button below the two alternatives. For the two alternatives in each trial, the one simulated with the same stiffness parameter as the target was denoted as the “matching”, and the other, with a different stiffness parameter, was denoted as the “distractor”. The three videos in each trial were always from different scenarios, and the mass of each cloth was randomly sampled from the five levels. The total number of unique trials was 480. For each observer, we randomly selected 100 trials by balancing task difficulty, 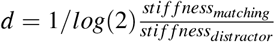, such that there were equal number of trails (i.e., 5) at each difficulty level.

## Computational model: cloth perception as mental physics simulation

We hypothesize that visually estimating the properties of soft objects in a dynamic scene amounts to probabilistic inference in a generative model that describes the mechanics and statespace of how soft materials respond to external forces applied to them. First, we describe the generative model to capture, in engineering-terms, our practical knowledge of how soft material works. Second, we describe an approximate inference procedure to condition this generative model on the observed object state and to make inferences about its underlying stiffness and mass properties.

### Generative model

The generative model consists of prior distributions over the physical properties (mass and stiffness) of a cloth and then simulating how it moves under the external forces in a scene. To capture the full variation of stiffness and mass observed in our task, we place uniform priors on mass and stiffness during scene initialization (mass *m*_0_; stiffness *s*_0_): *m*_0_ ~ Uniform(2^−2^,2^2^) and *s*_0_ ~Uniform(2^−7^,2^1^).

We implement the simulation process using a particle-based physics engine, FLeX, denoted Ψ. In FLeX, soft object mechanics, rigid body simulation, as well their interactions are all formalized as particle-particle interaction rules. Thus, it is referred to as a “universal” simulation engine as it can use the same substrate to simulate different forms of the matter. A cloth is simulated as a grid of particles connected to each other by massless springs that induce the effect of stiffness in that object.

To initialize a cloth simulation, we sample its physical properties (mass and stiffness), a scene configuration including shape and position of stationary physical objects (colliders) and moveable physical objects (e.g., feather, box), and any external forces (e.g., wind; gravity, which is turned on in all scenes; see Figure 3 “initialization”). The scene configurations are treated categorically, consisting of one of the four scenarios in our task. Once a scene is initialized, its statespace is unfolded over consecutive time steps, outputting simulated velocities (*v_i,j,t_* ∈ R^3^) and positions (*p_i,j,t_* ∈ R^3^) for each simulated particle *i, j* at each time step *t* (Figure 3). To avoid unnecessary clutter, we denote velocities and positions of all particles at time *t* as *v_t_* and *p_t_*.

**Figure 3:**
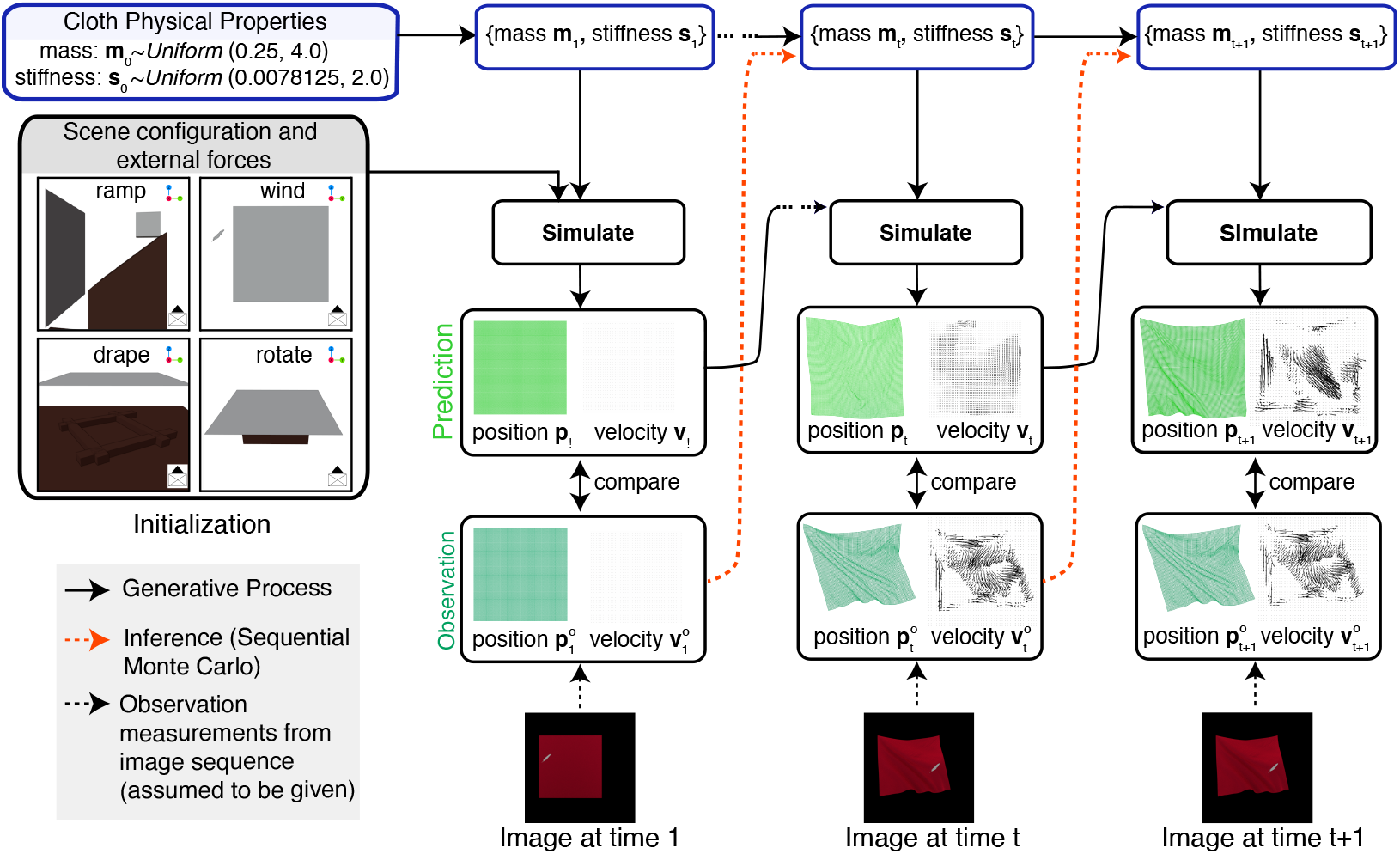
Schematic of our modeling framework. The generative model is described by the flowchart in the black frames and black arrows connecting them. This generative model takes as inputs a scenario (e.g., the “ramp” scenario), the external forces in the scene (e.g., gravity, wind), and the hidden physical properties of the cloth (mass *m_t_*; stiffness, *s_t_*), and simulates the cloth’s movements, predicting the cloth state measurements (*p_t_, v_t_*). This simulation process is implemented using a particlebased physics simulation engine (Macklin et al., 2014). We assume in the generative model that the 3D cloth state is observed (positions and velocities, 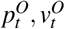 of each simulated cloth particle). We perform online belief updating in this model over video input using sequential Monte Carlo (SMC; inference conditionals are illustrated by the orange dashed arrows). At a given time step *t*, SMC compares the predicted state based on the simulated cloth (*p_t_, v_t_*) with those in the observation 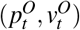, and accordingly updates its mass and stiffness beliefs.

Finally, in the generative model, we assume that the mass and stiffness values come from a temporal kernel such that they can change over time according to the following Gaussian distributions: *m_t_* ~ *N*(*m*_*t*−1_, *σ_m_*) and *s_t_* ~ *N*(*s*_*t*−1_, *σ_s_*). In the context of our work, we largely made this assumption for computational convenience; however, it is worth exploring the relationship between the perceptual persistence (Scholl, 2007) of soft objects and varying their mass and stiffness on the fly.

These conditional distributions induce the following posterior that we wish to estimate.

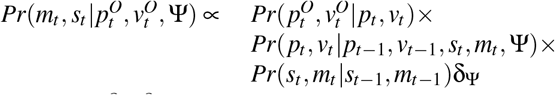

where 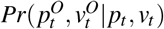 is the likelihood function (a normal distribution centered around the predicted positions and velocities with an observation noise of *σ*_O_) and *δ*_Ψ_ is a dirac-delta function at Ψ.

### Approximate inference using SMC

We hypothesize that perception of soft object properties can be cast as probabilistic inference in this simulation-based generative model. As an instantiation of our hypothesis, we aim to extract the mass and stiffness of the cloth, the material properties that causally influence the cloth’s behavior (shown as dashed orange arrows in Figure 3), while it is swivelling (rotate), waving in the wind (wind), falling on an uneven surface (drape) or being hit by an object (ramp).

We use the Sequential Monte Carlo (SMC) method to infer these latent parameters of a cloth given the observed state measurements 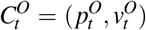, where 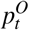 and 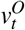 denote the observed positions and velocities of the simulated particles of the cloth at time *t*. Instead of computing these observations from an image, in this work, we assume that they are given. We come back to this assumption in the Discussion section. Overall, the goal of this inference procedure is to recover the posterior given by *Pr*(*m,s*|*C^O^*) = *P*(*C^O^*|*m,s*)*P*(*m,s*).

In our model, *N* SMC particles (as in particle filtering, not to be confused with the FLeX particles) are initialized with uniformly sampled mass *m*_0_ and stiffness *s*_0_ values. Each particle passes their latents and state into the FLeX engine to generate the next predicted cloth states {*C*_1_}_N_ = {(*p*_1_, *v*_1_)}_*N*_. SMC computes the score for each particle by comparing {*C*_1_}*N* with the observed cloth state 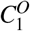 and randomly samples *N* new particles in the proportion to their scores. We implemented this inference procedure using a state-of-the-art probabilistic programming package called Gen (Cusumano-Towner, Saad, Lew, & Mansinghka, 2019).

### Simulation details

In all our experiments, we use a cloth with 105 x 105 grid of vertices, with homogeneous mass and spring stiffness values across all cloth particles (FleX particles).

In our SMC simulations, we use 20 particles in all four scenarios. We assign the observation noise *σ_O_* = 1.0. The kernel variance parameters for mass and stiffness are as follows:

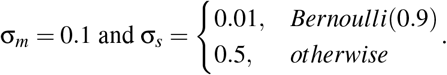

For each input observation, we simulate 7 SMC chains. In our behavioral comparisons, we treat each of these chains as a simulated subject and report their average performance.

## Behavioral experiment

### Participants

We collected data from 18 participants using Prolific, a crowdsourcing platform. All participants were naive to the purpose of the experiments. They agreed with an electronic informed consent prior to the experiment, and received a compensation of $4.75 for their participation. This study was approved by the Yale University Institution Review Board (IRB). One participant failed to meet the inclusion criteria (described in the Results below) and thus was excluded.

### Procedure

The experiment was conducted using PsiTurk (Gureckis et al., 2016). At the beginning of the experiment, the participants were provided with a written definition of cloth stiffness, an explanation of the task, a description of the four scenarios, and a few example animations illustrating these scenarios and the most extreme physical parameter pairings (e.g., softest vs. most stiff and heaviest vs. lightest cloths; 8 animations in total). At the end of these instructions, participants were presented with three quiz questions measuring their comprehension of the task and stimuli. The participants needed to answer all three questions correctly to proceed to the main experiment. Otherwise, they were directed back to the start of the instructions.

On each trial, participants were presented with video triads (Figure 2B) and they were asked to perform a 2AFC stiffness matching task (described in Section Task). The response region was hidden until the participants watched each video stimuli for 5 sec. Participant indicated their choice by clicking the “left” or “right” button beneath the alternative videos, and they could change their response of the current trial by clicking the other response button. There was no time limit for each trial and participants was instructed to click the “Next” button on the bottom center of the experimental page to move on to the next trial. Each participant completed 100 unique trials.

## Results

As an inclusion criteria, we calculated the accuracy (*ACC*) of each participant’s matching performance on the 60 easiest trials (|*log*_2_(*s_matching_*) − *log*_2_(*s_distractor_*)| = 4, 6, 8, respectively). One participant (*ACC* = 0.57) failed to perform better than chance level on these trials, and were excluded from the experiment. The following analysis was conducted on data from the remaining 17 participants (*ACC* = 0.74 on these easiest 60 trials, *SD* = 0.005).

The left panel in Figure 4B(1) shows the accuracy pooled across all participants for each possible stiffness pair between the matching and distractor item. Each “blue grid” shows a stiffness pair; the accuracies reported in these blue grids were averaged across different mass and scenarios. Overall, participant’ accuracy was reasonably high considering the difficulty of this task, *ACC* = 0.67, *p* < .001. We observed that as the stiffness values of the matching and distractor cloths became more similar, participants had a harder time to differentiate between them, as is suggested by a decrease in the matching accuracy. For example, for the pair of a very soft matching cloth (2^−7^) and a very stiff distractor cloth (2^1^), participants’ matching accuracy was 0. 91; and the accuracy dropped to 0.53 when the distractor cloth had a similar stiffness value (2^−5^) to that of the matching cloth (2^−7^).

**Figure 4:**
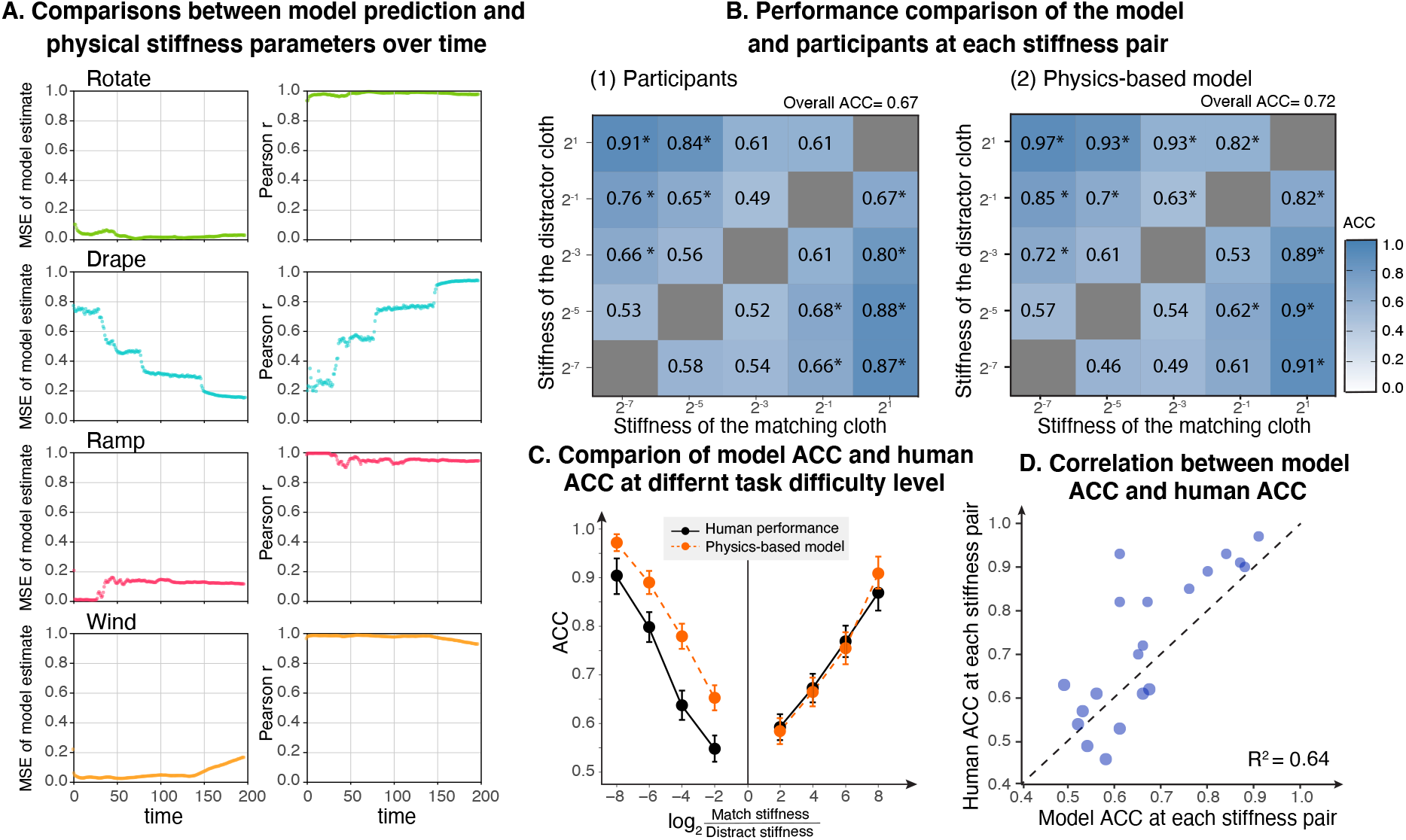
Overview of the behavioral and modeling results. (A) The model’s stiffness estimates become more accurate over the course of the 200 video frames that it observes. (B) We present the performance of the humans (left) and models (right) by aggregating the data with respect to the stiffness values of the matching and distractor items. We have a total of 20 such pairs (each “blue grid”). (C) Performance of the model and behavior as a function of the difference (or difficulty) between the stiffness values of the matching and distractor items. (D) Correlating the two panels from (B) to quantitatively evaluate the relationship between the model’s and humans’ accuracy at each stiffness pair.

Next, we measured the behavioral accuracy as a function of the stiffness difference between the matching and distractor cloths (Figure 4C, black line). We found the matching performance dropped as the stiffness difference became smaller in a log-linear fashion, which was consistent with previous findings (Bi et al., 2018). Last, we sought to test whether the differences in mass values impacted participants’ stiffness matching performance. To to do, we fitted a linear regression model to predict stiffness matching accuracy at each difficulty level (i.e., *ACC* reported in each grid of Figure 4B(1)), using stiffness difference (*S_diff_* = *s_matching_* – *s_distractor_*) and categorical mass difference

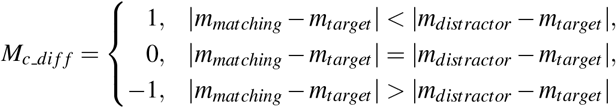

The model was fitted as below: *ACC* = 0.10 × *S_diff_* + 0.007 × *M_c_diff_* + 0.003 × *S_diff_* × *M_c_diff_* + 0.46. The regression model explained significant variance in the responses, *F*(3,1696) = 1438.0, *p*< .001, *R*^2^ = .72. The regression coefficient for the stiffness difference term (i.e., *S_diff_*) was significant, *t*= 63.6, *p*< .001, suggesting that the stiffness difference between the target and matching significantly impacts the behavioral accuracy. Additionally, the coefficient for the mass difference term (i.e., *M_c_diff_*) was marginally significant, *t*= 1.721, *p*= .08, suggesting that the mass has a marginal effect on stiffness perception.

Overall, the behavioral results suggest that humans are able to generalize across different scenarios and match cloth stiffness substantially above chance levels. This type of generalization is suggestive of physics-based internal representations of soft objects; In the next section, we turn to computational modeling to further test our hypothesis, by making comparisons to behavior.

## Testing models as accounts of human behavior Simulation results

Figure 4A shows the accuracy of our model’s inferences across the four scenarios (rotate, drape, ramp and wind). The model’s stiffness estimates generally become more accurate as the model accumulates evidence over the course of 200 frames (Figure 4A, first column). Despite the distance to the ground truth values, the model’s stiffness inferences preserve the rank order of the stiffness values within a scenario (Figure 4A, second column).

### Comparisons to behavior

We now evaluate our model by making quantitative comparisons to behavior. We start with noting that the model’s average accuracy is a little bit higher than behavior (0.72 vs. 0.67).

Figure 4B shows finer grained comparisons between the model and behavior. We present the performance of the model across all stiffness pairings between the matching and distractor items in Figure 4B(2) (integrating over mass variations). We note that the patterns of accuracy in the model across the different difficulty levels (in terms of the difference between the matching and distractor stiffness values) are largely consistent with what we observe in behavior (Figure 4B(1)). We quantify the correspondence between these accuracy patterns of the model and humans in Figure 4D (*R*^2^ = 0.64).

To determine whether mass has any effect on this model’s performance, we apply the regression analysis, described in the behavioral section, to the performance of this model. We find no main effect of the mass (*p*>.05), however the interaction term between mass and stiffness was significant (*p*<.01) indicating that model’s estimates were impacted by its mass inferences.

Finally, we compare the model’s performance to behavior at the level of stiffness differences (Figure 4C). We find that like humans, the model is largely linear in log scale.

## Discussion

In this work, we consider the hypothesis that the soft material perception can be understood in terms of an abstract, physics-based representation of such soft objects. To evaluate this hypothesis, we presented a cloth perception experiment that required generalization across scenarios—to our knowledge this is the first experiment to test such generalization in the context of soft materials. We also presented a simulation-based computational model of cloth material perception, finding that inferences in this model can explain behavior.

Our work adds to the existing literature showing that humans’ intuitive physical scene understanding can be best understood in terms of an underlying mental simulation engine, to represent how objects move and interact (Battaglia et al., 2013; C. Bates, Battaglia, Yildirim, & Tenenbaum, 2015; C. J. Bates et al., 2019; Yildirim et al., 2020; Wu et al., 2015). We extend this line of work to the domain of soft object perception, visiting cloths as a starting point. We are excited to further explore the scope of this approach in cloths and in soft object perception more generally.

A core aspect of our proposal is that the brain might process soft materials in terms of non-rigid physical systems. Neurological studies using “minimal” dot stimuli that can elicit a variety of physical material perceptions, such as flowing liquids, crumbling cubes, flapping cloth, and bouncing cube (Schmid & Doerschner, 2018; Bi, Jin, Nienborg, & Xiao, 2019) find that such objects recruit a wide network of areas in the brain (Schmid, Boyaci, & Doerschner, 2020). Most of these regions seem to overlap with the “intuitive physics regions” reported in (Fischer, Mikhael, Tenenbaum, & Kanwisher, 2016). This correspondence is consistent with our perspective that soft material perception involves some sort of a physical simulation and we hope to explore this possibility in future studies.

